# 3D reconstruction of genomic regions from sparse interaction data

**DOI:** 10.1101/2020.10.11.334847

**Authors:** Julen Mendieta-Esteban, Marco Di Stefano, David Castillo, Irene Farabella, Marc A Marti-Renom

## Abstract

Chromosome Conformation Capture (3C) technologies measure the interaction frequency between pairs of chromatin regions within the nucleus in a cell or a population of cells. Some of these 3C technologies retrieve interactions involving non-contiguous sets of loci, resulting in sparse interaction matrices. One of such 3C technologies is Promoter Capture Hi-C (pcHi-C) that is tailored to probe only interactions involving gene promoters. As such, pcHi-C provides sparse interaction matrices that are suitable to characterise short- and long-range enhancer-promoter interactions. Here, we introduce a new method to reconstruct the chromatin structural (3D) organisation from sparse 3C-based datasets such as pcHi-C. Our method allows for data normalisation, detection of significant interactions, and reconstruction of the full 3D organisation of the genomic region despite of the data sparseness. Specifically, it produces reliable reconstructions, in line with the ones obtained from dense interaction matrices, with as low as the 2-3% of the data from the matrix. Furthermore, the method is sensitive enough to detect cell-type-specific 3D organisational features such as the formation of different networks of active gene communities.

## Introduction

Chromatin within the nucleus is organised into higher order structures that emerge at different genomic scales, from chromosome territories (at tens of megabases scale), active and inactive chromatin domains (at few megabases scale) [1], self-interacting domains or TADs (at hundreds of kilobases scale) [2-4], and long-range chromatin loops between regulatory elements (at tens of kilobases scale). This multi-scale organization has a direct impact on many biological processes such as gene regulation, DNA replication, and cell differentiation [5-7]. Indeed, genome structure typically reflects cell-type-specific differences in the transcription pattern, and it is frequently rewired upon cell state changes and disease onset [8]. Thus, investigating the principles shaping chromosome three-dimensional (3D) structure is pivotal to shed light into the relationship between genome structure and function.

Several experimental techniques are available to examine chromatin organisation [9]. Among them, molecular biology methods such as Chromatin Conformation Capture (3C) and its derivatives are widely used [10]. These experiments retrieve information about the frequency of interaction between loci in single [11-13] or in populations of thousands to millions of cells and have been designed to analyse the chromatin landscape at different genomic scales [1, 14-16]. For example, some cell population-based experiments allow the retrieval of unspecified interactions in the whole genome (*e.g*., Hi-C [1], Micro-C [14], GAM [15], and SPRITE [16]). Complementarily, other 3C-based experiments are tailored to capture interactions centred on a specific locus with the rest of the genome (e.g., 4C [17] and multi-contact 4C (MC-4C) [18]) or on sets of dispersed loci in the genome, such as loci enriched for a specific protein (HiChIP) [19] or loci harbouring gene promoters (pcHi-C) [20]. Each class of 3C-based experiments provide different but complementary insights on particular aspects of the genome organization, and their analysis is dependent on the experimental genomic resolution and on the inherent technical biases of each experimental procedures.

A variety of physics- and data-driven approaches for genome 3D reconstruction have been developed to expose the principles shaping chromosome 3D structure [21-24]. For instance, data-driven (restraint-based) modelling approaches as PSG [25, 26], TADbit [27], 4Cin [28], and TADdyn [29] have been implemented to reconstruct ensembles of chromatin 3D models from cell population-based datasets. Others are focused on the 3D modelling of chromatin based on single-cell Hi-C data, like manifold based optimization [30] and NucDynamics [31]. However, the majority of the data-driven methods are based on interaction experiments that have been designed to retrieve dense contact information from a continuous set of loci or the whole genome, while other interaction experiments are characterised by data sparseness (*e.g*., HiChIP or pcHi-C). As such, data-driven methods for sparse data modelling are needed.

Generally, the interaction profiles of sparse 3C-based datasets have specific properties that set them apart from other 3C-like techniques characterised by a dense interaction profile. Indeed, protein or promoter capture-based interaction profiles are heavily biased on interactions between captured fragments and devoid of interactions between non-captured fragments. This fact poses the question of whether this lack of information prevents the 3D reconstruction of the whole loci of interest and its analysis, or whether it is sufficient to allow for accurate 3D modelling. To answer this question, we have implemented a new method, which is tailored to integrative modelling and analysis of sparse 3C-based datasets. We have also validated the procedure comparing the resulting reconstructed models with available dense experimental datasets, unveiling that the 3D chromatin organisation can be well recovered by interrogating only a small percentage of loci. Additionally, we have designed new tools to facilitate a robust differential analysis of the resulting models and showcased their usability in comparative analyses using the β-globin locus as a test case. Interestingly, comparing different cell-types, we unveiled that the β-globin locus in cord-blood Erythroblasts (cb-Ery), where its foetal and adult β-globin genes are highly expressed, is hierarchically organised in a 3D network of active gene communities that follows an expression gradient.

## Results

### Overall modelling strategy for sparse 3C data

Sparse 3C datasets provide information of interactions that involve a limited number of specific loci in the genome. pcHi-C, for example, provides a promoter-centred view of chromatin interactions, helping to assign distal regulatory regions to their target genes, thus providing insights on how gene expression might be controlled [32-34] and how disease-associated genomic variation could affect gene regulation [35]. The main limitation of these sparse technologies, however, is the scarcity of specialized tools for their analysis. Here we have developed an integrative 3D modelling method capable of dealing with data sparsity, enabling the analysis and interpretation of pcHi-C data, and tested it on 12 distinct loci (Benchmarking datasets; **Methods** and **Supplementary Table 1)**. Our method follows an integrative modelling procedure comprising five steps [36]: (i) gather experimental data and process them to obtain the input interaction matrix for the modelling approach, (ii) represent the selected chromatin regions using a bead-spring polymer model with a particle size proportional to the genomic resolution of the experimental data, (iii) transform the frequency of interactions into spatial retrains, (iv) sample the conformational space by steered molecular dynamics, and (v) analyse and validate the obtained ensemble of 3D models (**Methods** and **Figure 1A**).

**Figure 1.**
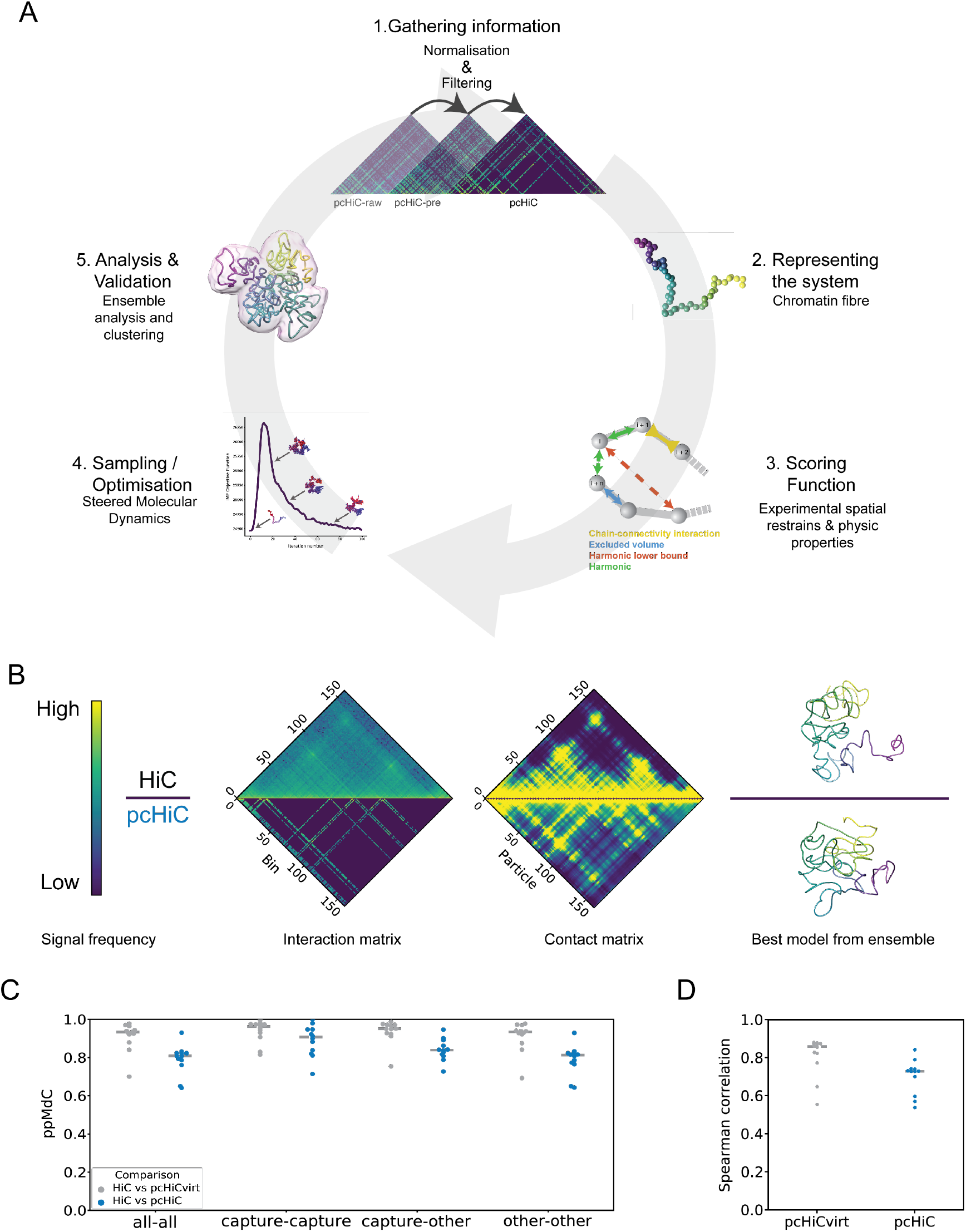
Integrative modelling for sparse datasets efficiently reconstructs the 3D organisation of genomic loci. **(A)** Workflow of the integrative modelling approach followed to build ensembles of chromatin 3D models from pcHi-C: i) gathering the input interaction matrices with subsequent normalisation and filtering; ii) representation of the chromatin fibre as a polymer with the particle size proportional to the resolution of the experiment; iii) definition of the scoring function used in the modelling procedure. Here, the scoring function comprises spatial restrains derived directly from the input interaction data and from properties of the chromatin fibre (**Method**); iv) sampling the conformational space by steered molecular dynamics (**Method**); and v) validation of the obtained ensemble of models and further analysis. Model images in all panels were created with Chimera [73]. **(B)** Representation of the input and output data from region 2 (**Supplementary table 1**). The upper half of the panel refer to the dense dataset (Hi-C), whereas the lower half refer to the sparse-datasets (pcHi-C). From left to right, the matrices of normalised interaction frequency (**Methods**) between each pair of bins, the contact matrix obtained from the ensemble of models of region 2 displays the percentage of models in which two bins are found bellow the defined distance cut-off for the contact (**Methods**), and the best model from the ensemble as assesses by the scoring function. The colour bar shows the colour coding from low (blue) to high (yellow) interaction or contact frequencies signal. (**C)** Comparison between models ensembles derived from sparse (pcHi-Cvirt and pcHi-C in grey and blue, respectively) and dense (Hi-C) datasets assessed by the particle-to-particle median distance correlation (ppMdC; **Methods**). Three subsets of particles have been compared given the enclosed loci: (i) captured loci (capture), (ii) non-captured loci (other), and (iii) all the loci (all). The grey dashed line indicates the median ppMdC in the 12 analysed regions. **(D)** Element-wise Spearman correlation coefficients between the experimental Hi-C interaction matrices and the contact maps derived from the model ensembles reconstructed from sparse data (pcHi-Cvirt and pcHi-C in grey and blue, respectively). The grey dashed line indicates the median element-wise Spearman correlation coefficients in the 12 regions analysed.

In this work, we gathered pcHi-C interaction data (**Methods**), whose processing step is pivotal to minimize the experimental biases from the capture protocol. To this end, we designed a multi-stage normalisation procedure named PRoportion of INTeraction approach (PRINT, **Methods**). PRINT weighs each interaction by dividing it by the cumulative whole-genome interaction frequencies of both of the interacting bins, regularising the interaction patterns for the fact that captured loci are highly enriched in contacts. It also removes the pcHi-C unspecific interactions between non-probed bins. To test quantitatively the performance of our normalisation procedure, we compared each of the normalisation stages of the pcHi-C matrices with the respective Hi-C matrices normalised with OneD in each of the selected loci [37]. The median correlation between bins with interaction data in both matrices was 0.27 (+/- 0.025 Median Absolute Deviation (MAD)) for raw pcHi-C matrices (pcHi-C-raw), increasing to 0.44 (+/- 0.032 MAD) with the pcHi-C pre-normalisation step (pcHi-C-pre), and reaching 0.60 (+/- 0.056 MAD) for fully normalised pcHi-C matrices (pcHi-C-norm) (**Supplementary Figure 1A**), suggesting that PRINT reduced successfully the target biases. Then, we represented the selected loci as a bead-spring polymer model with a particle size set to 5 kb, taking into account the restriction fragment lengths distribution in the benchmarking datasets (**Supplementary Figure 1B**). Similarly to TADbit [27] and TADdyn [29], to simulate the structural conformation of genomic loci, we then transformed the interaction frequencies associated with each bin pair into spatial restraints (**Methods**). The latter were then imposed on the model using steered molecular dynamics as sampling method in which the spring constant associated to each restraint was ramped up as a function of simulation time from zero to the value computed from the interaction data. Lastly, we implemented new means for a robust quantitative spatial differential analysis of genomic loci.

### Comparison between sparse and dense 3C-derived models

Dense Chromatin Conformation Capture data has been extensively used to reconstruct the 3D organisation of genomic loci [25, 27, 29, 30]. Here, to test the reliability of our modelling approach, we used sparse and dense datasets to build ensembles of models of the same loci. Specifically, we applied our integrative method for sparse data modelling to previously published pcHi-C datasets of GM12878 cells [32] to reconstruct 3D model ensembles of 12 distinct loci (**Figure 1B** and **Supplementary Table 1**) at a 5kb resolution and compared them with the corresponding ones reconstructed using Hi-C [6] at the same genomic resolution. Additionally, to quantify the effect of sparsity in the comparison independently of the experimental protocol biases, we generated virtual pcHi-C (pcHi-Cvirt) interaction matrices from the normalised Hi-C datasets extracting the rows and columns probed in the pcHi-C experiment (**Methods**). These virtual sparse matrices were then used to reconstruct 3D model ensembles of the selected loci.

The comparison between the sparse and dense derived 3D model ensembles revealed that it is possible to recover most of the 3D organisation of the dense dataset in spite of the data sparsity (**Figure 1C**). Indeed, the all-vs-all particle-to-particle median distance correlation (ppMdC) between the sparse and dense derived 3D model ensembles was 0.81 (+/- 0.019 MAD) and 0.93 (+/- 0.024 MAD) for both pcHi-C and pcHi-Cvirt. Additionally, when comparing distances between particles that have both been captured in the pcHi-C experiment (capture-capture), the ppMdC was higher, reaching 0.91 (+/- 0.054 MAD) for pcHi-C and 0.96 (+/- 0.019 MAD) for pcHi-Cvirt. Consistently, when comparing distances between non-captured particles with captured particles (capture-other) or between non-captured particles (other-other), the ppMdC indicated good agreement with values of 0.84 (+/- 0.03 MAD) and 0.95 (+/- 0.02 MAD), and 0.81 (+/- 0.02 MAD) and 0.93 (+/- 0.02 MAD) respectively for pcHi-C and pcHi-Cvirt in both comparisons (**Figure 1C**). The results indicate that the sparse derived ensembles of 3D models are a good representation of the dense experiment and that the intrinsic experimental biases of the capture experiment only minorly affect the 3D reconstruction. Indeed, comparing the whole contact map computed from the 3D model ensembles derived from sparse data directly with the whole experimental Hi-C interaction matrices revealed that the reconstructed ensembles of models are in good agreement with the dense experimental data having an element-wise Spearman’s rank correlation coefficient of 0.73 (+/- 0.02 MAD) and 0.86 (+/- 0.02 MAD), for pcHi-C and pcHi-Cvirt derived ensembles of models, respectively (**Figure 1D**). Overall, this suggest that the ensembles of models reconstructed by our approach represent well the 3D organisation of the selected genomic regions and, more importantly, recover the spatial arrangements of loci that are not interrogated by the sparse experiment.

### Reconstruction efficiency and data sparsity

To investigate the relationship between the reconstruction efficiency and data sparsity, we simulated ‘synthetic’ capture data. Briefly, we generated 10 different sets of ‘synthetic’ capture matrices that represent generic capture-like experiments. We started from the contact matrix derived from a 3D toy-genome models ensemble that simulates roughly a one Mb length genome (comprising more than 600 particles) with a TAD-like architecture, a high level of interaction noise, and low variability between models [38] (**Methods** and **Figure 2A**). To build each of the 10 ‘synthetic’ sets, we randomly selected 22 captured loci and constructed 6 additional datasets of different sparsity down-sampling each set considering 2, 4, 6, 10, 14, and 18 loci at a time, which mimics the distribution of captured probes per Mb present in a typical genome-wide pcHi-C experiment (**Figure 2B**). The constructed 70 capture-like matrices thus aim to represent typical pcHi-C experimental design. Using our integrative modelling method for sparse datasets, we reconstructed, from each of the ‘synthetic’ capture matrices in the dataset and their down-sampled counterparts, ensembles of 100 models, and compared them with the reference toy-genome ensemble (**Figure 2A**). Independently of the sets, the ppMdC between the sparse and dense model ensembles increased with the number of captured particles used in the modelling procedure reaching a median correlation between sets of 0.82 (+/- 0.02 MAD with just 10 captures per Mb (**Figure 2C**). Notably, also with 4 and 6 captures per Mb the ppMdC reached 0.69 (+/- 0.04 MAD) and 0.79 (+/-0.05 MAD) for 4 and 6 captures, respectively, although with greater variation within sets. This suggests that with 10 captured loci per Mb the uncertainty in the input information is smaller, leading to more precisely reconstructed models. Nevertheless, it is possible to reconstruct good models also with fewer as 4 captured loci per Mb although with a higher degree of variability. To quantify the effect of data sparseness on model reconstruction, we next measured the amount of input information used during the modelling as the percentage of all possible interaction pairs in the contact matrix (dense data input) and then assessed it with the ppMdC. The results indicate that it was possible for the majority of the sets (8/10) to reliably reconstruct the reference toy genome (ppMdC > 0.8) with just 2-3% of all the interaction pairs in the contact matrix used as restrains (**Figure 2D**). Taken together, this analysis shows that it is possible to consistently recover most of the 3D organisation of a region of interest with 10 captured loci per Mb and with just 2-3% of all possible interactions within a region captured.

**Figure 2.**
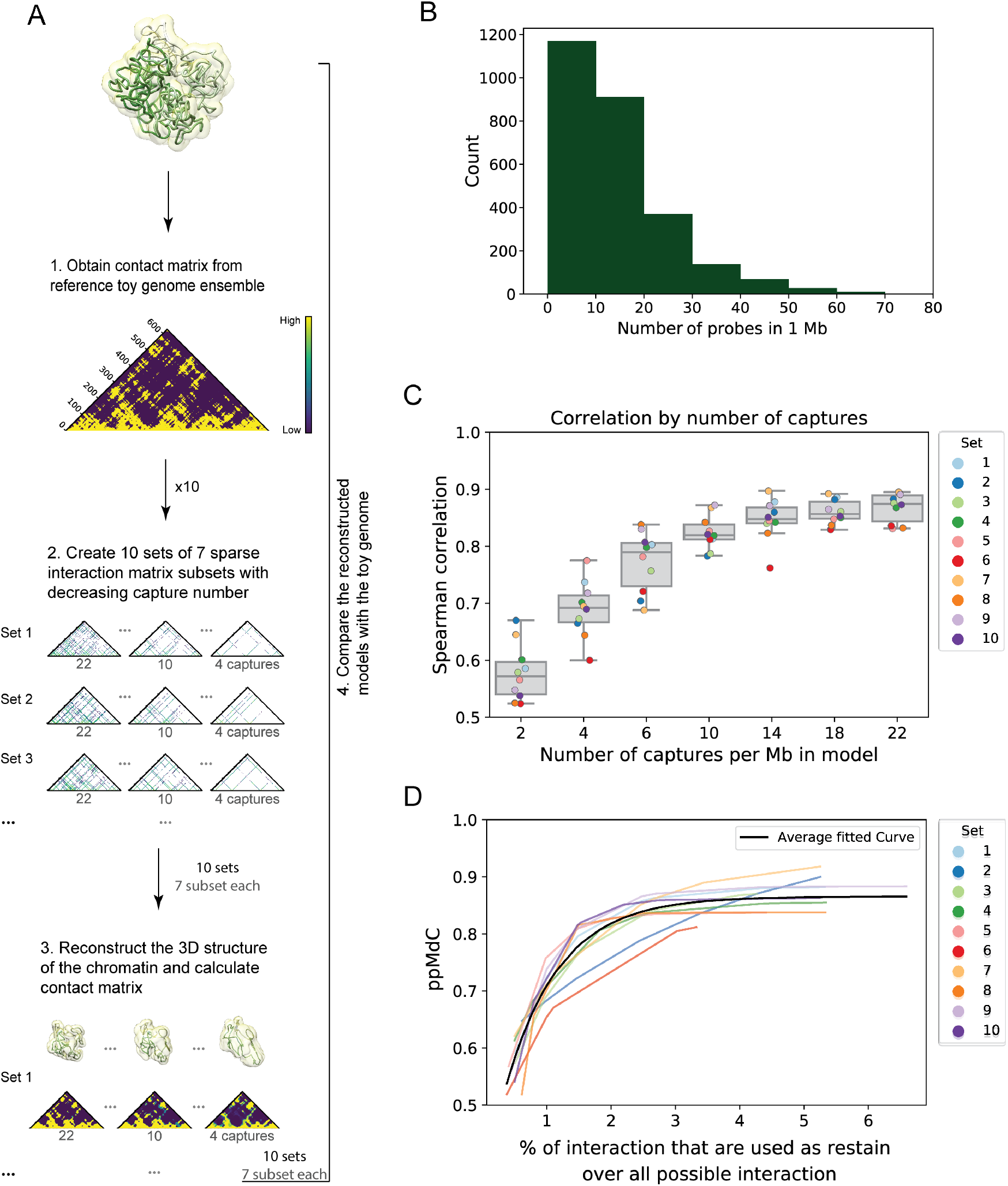
A low percentage of the interaction data is needed to produce reliable 3D reconstructions. **(A)** Workflow for the generation of 3D model ensembles from ‘synthetic’ sparse datasets and comparison with the toy genome. A total of 70 ‘synthetic’ captured map were generated representing 10 different capture experiments with different level of data sparsity (**Methods**). Model images were created with Chimera [73]. **(B)** Distribution of pcHi-C probes per megabase windows in the genome [32]. **(C)** Distribution of the ppMdC between the ‘synthetic’ models and the toy genome grouped by subsets of captures per megabase. Box boundaries represent 1st and 3rd quartiles, middle line represents median, and whiskers extend to 1.5 times the interquartile range. The ten sets of captured positions are displayed with the colour code shown in the insert. **(D)** Relationship between the ppMdC and the percentage of cells in the matrix used as restrains in each set represented with an exponential fit. The used colour code is the same as in **C**, the grey line represents the mean fit of all the datasets in analysis.

### Cell-type specific organisation of the β-globin locus

To illustrate the utility of our integrative approach in unveiling the differential organisation of loci, we applied it to the genomic region surrounding the β-globin locus in 3 different cell types for which pcHi-C data are available [33], namely cord-blood derived Erythroblasts (cb-Ery), naive CD4+ T-cells (nCD4), and Monocytes (Mon). The selected genomic region contains five coding genes (HBB, HBD, HBG1, HBG2, and HBE1) with developmental-stage-dependent expression [39], which is finely regulated by a set of upstream enhancers known as the Locus Control Region (LCR) [40]. This locus is known to be in an active conformation in cb-Ery, where the LCR is interacting mainly with expressed genes as HBB and HBD, but not in nCD4 and Mon cells [33].

First, we defined the optimal region to be modelled based on the interaction networks (in all cell types) of the embryonic (HBG1 and HBG2) and adult (HBB and HBD) globin genes with the rest of the genome at 5 kb resolution (**Methods**). The defined region spanned 4.7 Mb of chr11 (chr11:3,795,000-8,505,000 base-pairs (bp)) comprising several neighbouring genes and multiple long-range regulatory elements. By applying our integrative approach, we generated an ensemble of 1,000 3D models for each cell type. The packing of the genomic region was significantly different in each cell types with median radius of gyration of 248+/-3, 242+/-2, and 237+/-2 nm for cb-Ery, nCD4 and Mon, respectively (p-values < 9.1e^-163^ in each of the pairwise comparisons using two-samples Kolmogorov-Smirnov statistics) (**Supplementary Figure 3A**), with the topology of the region in cb-Ery being less tightly packed than in nCD4 and Mon. Each ensemble was then clustered by structural similarity [27] and the models from the most populated cluster were selected for the comparative analysis between cell-types. Clustering by distance root-mean-square deviation (dRMSD), confirmed that the topology of the region was markedly different in the three cell types, with nCD4 and Mon folds being more similar between each other than with cb-Ery (**Figure 3A**). Particularly interesting is how the topology of the β-globin locus (chr11:5,201,270-5,302,470) varied in the three cell types. Indeed, in Erythroblasts the β-globin locus appeared to be located further from the main core of the region as compared with naïve CD4+ T-cells and Monocytes, with median distances between the centre of mass of the β-globin locus of 286, 243, and 207 nm in cb-Ery, nCD4, and Mon, respectively (p-values < 3.46e^-101^ in all the pairwise cell-type comparisons; two-samples Kolmogorov-Smirnov statistic) (**Supplementary Figure 3B**).

**Figure 3.**
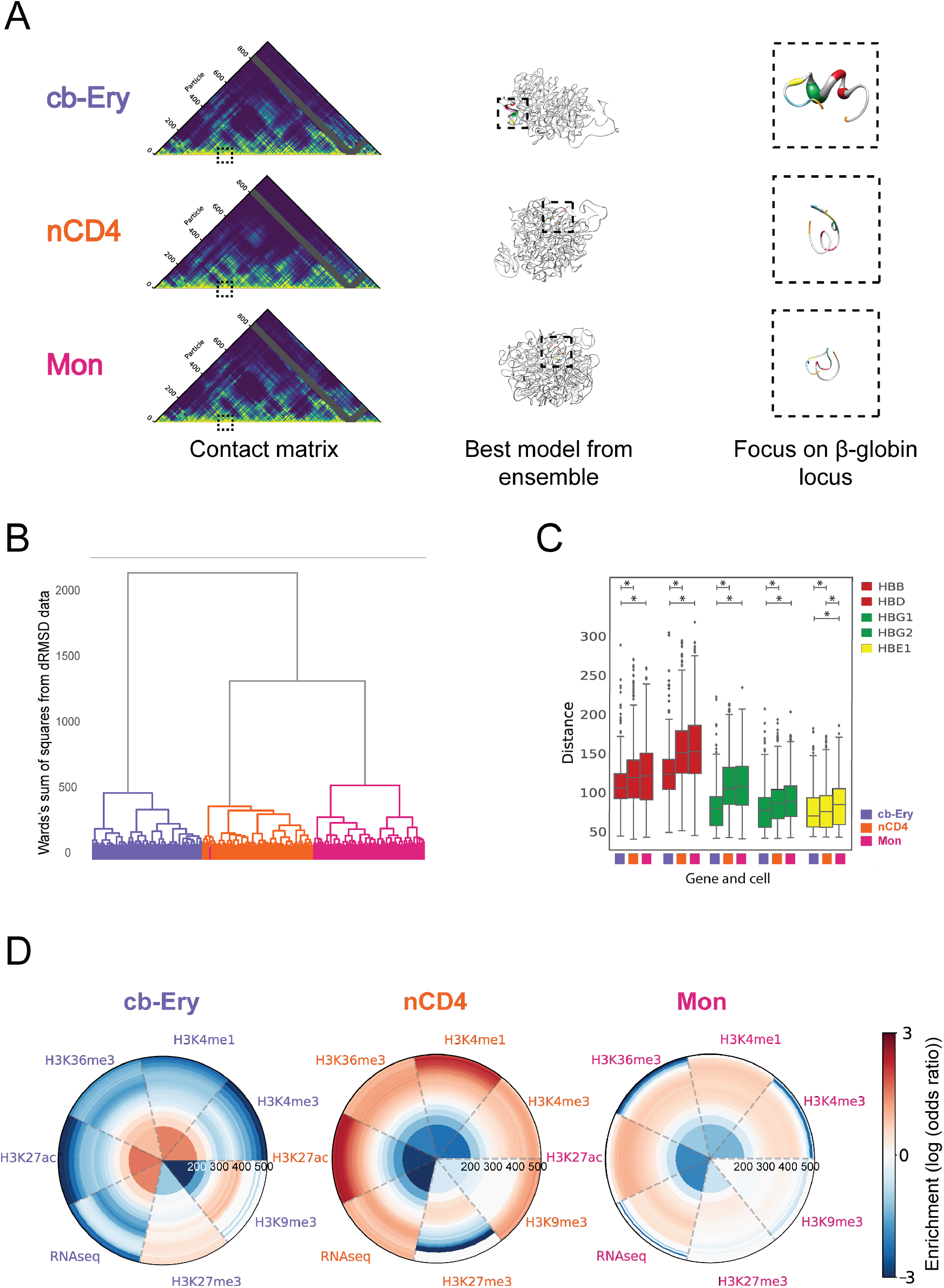
Cell-type specific organisation patterns of the β-globin locus. **(A)** β-globin locus in cb-Ery, nCD4, and Mon cell-types. From left to right: representation of the contact matrix derived from each of the model ensembles colour coded from low (blue) to high (yellow) contact frequency (columns filtered due to low interaction data are coloured grey); best model from ensemble as assesses by the scoring function; zoom up of the β-globin locus in the model. Models are represented as a tube with thickness proportional to the cell-type expression profile (**Methods**), the regulatory elements and genes in the β-globin locus are coloured as follow: HBB and HBD in red, HBG1 and HBG2 in green, HBE1 in yellow, LCR in blue and 3’HS1 and HS5 in orange. Model images were created with Chimera [73]. **(B)** Clustering tree (see *Hierarchical clustering of ensembles of 3D models* in Chromatin ensemble 3D analysis) of cb-Ery (purple), nCD4 (orange) and Mon (pink) model ensembles. **(C)** Cell-type specific distance distributions between the particle containing HS3 site of the LCR and the β-globin genes (HBB, HBD, HBG1, HBG2, and HBE1, colour coded as in (A)) as observed in the ensemble of models. Box boundaries represent 1st and 3rd quartiles, middle line represents median, and whiskers extend to 1.5 times the interquartile range (two-samples Kolmogorov-Smirnov test, asterisk indicate p < 0.007). **(D)** Radial plot showing the 3D enrichment around HS3 (**Method**). Each circumference shows the enrichment or depletion of features around HS3 on layers (up to 560 nm away from HS3) of non-overlapping volumes equal to the one of the initial sphere with radius of 200 nm. The colour bar shows the colour coding from highly depleted (blue) to highly enriched (red) features.

To characterise this further, we focused specifically on the β-globin locus and quantified its spatial organisation with respect to hypersensitive site 3 (HS3) in the LCR, which is forming an intricate network of interaction with the β-globin genes [41] and is required for their activation [42]. In line with this evidence, in the 3D ensemble of models representing cb-Ery cells, HS3 was significantly closer to HBB, HBD, HBG1, HBG2, and HBE1 genes than in the 3D ensemble of models representing nCD4 and Mon (p-values < 0.007, two-samples Kolmogorov-Smirnov test). In the latter two cell-types HS3 had a similar distance distribution with HBB, HBD, HBG1, and HBG2 genes (p-values > 0.01, two-samples Kolmogorov-Smirnov test) (**Figure 3B**).

Performing 3D enrichment analysis of varied epigenetic features and expression levels around HS3 (**Methods**), we unveiled a stark enrichment of active chromatin marks (H3K27ac, H3K36me, H3K4me1, and H3K4me3) and expression levels, and a clear depletion of inactive marks (H3K9me3 and H3K27me3) in cb-Ery. This 3D functional signature was absent in nCD4 and Mon, where active chromatin marks and transcript levels were depleted (**Figure 3C**). Overall, our models recapitulated the different 3D organisation of the β-globin locus and highlight the existence of a specific 3D functional signature enriched in active chromatin features that characterised the active β-globin locus in cb-Ery.

### Active gene communities in cb-Ery: a cell-type specific 3D signature

To examine whether the specific 3D functional signature of the active β-globin locus influence its genomic neighbourhood, we investigated its long-range interaction patterns. Comparative analysis of the distance profile between HBG2 (the most expressed gene in cb-Ery) and each of the selected loci (chr11: 3,795,000-8,505,000 bp), revealed the existence of an intricate cell-type specific network of spatially proximal expressed genes (**Figure 4A**), in line with previous observations of transcribed genes co-localizing in space [24, 43-46]. This network comprised distal transcribed sites (even located at 1.4 Mb away as STIM1) that showed cell-type specific spatial proximity. Indeed, HBG2 in cb-Ery was in closer proximity with all other expressed loci of the genomic neighbourhood than in nCD4 and Mon (**Figure 4B**).

**Figure 4.**
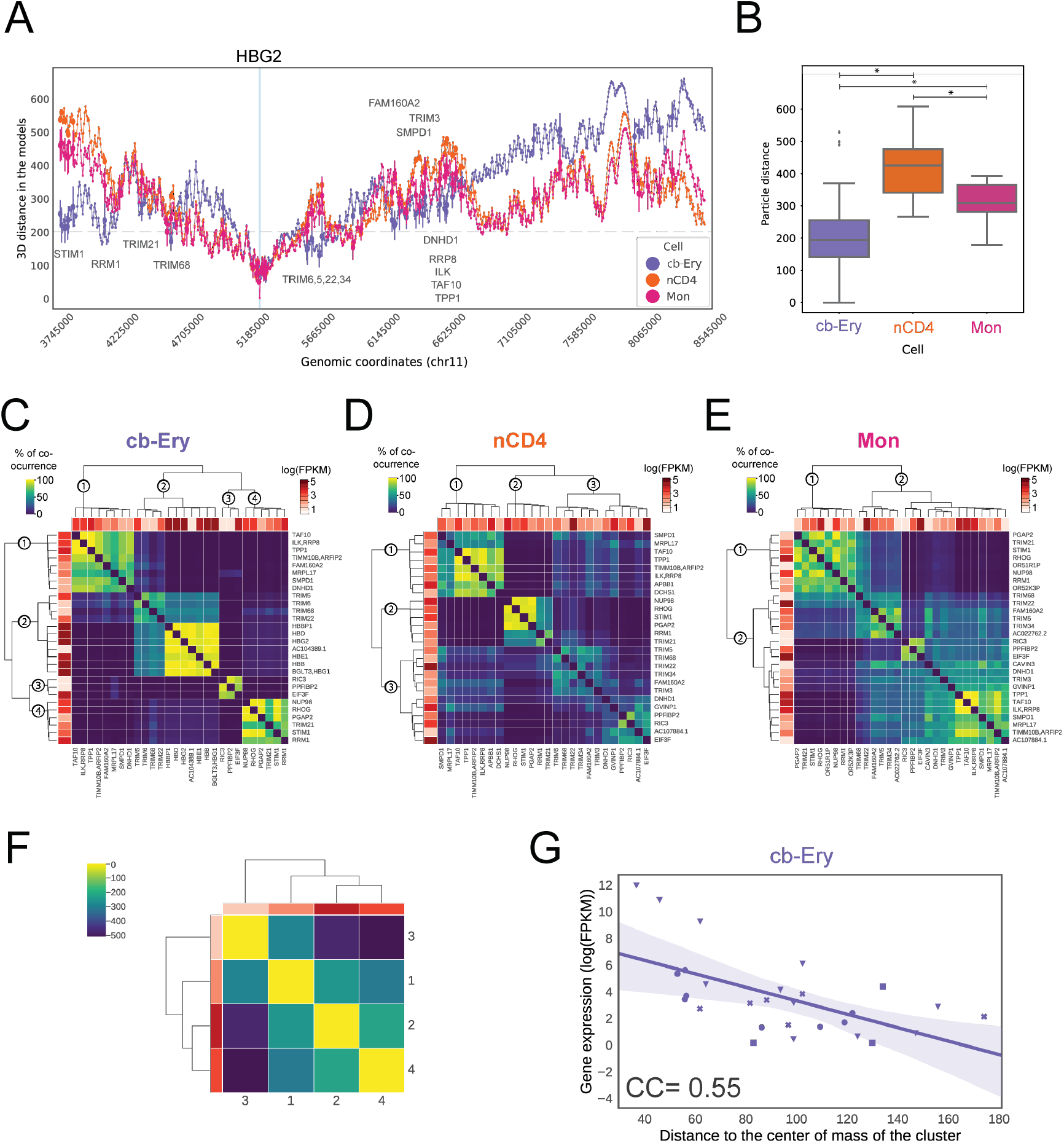
Communities of active genes as a cell-type specific 3D signature in cb-Ery. **(A)** Line plot of the mean distances between the TSS of HBG2 (focus point, blue vertical line) and all other particles in the genomic region (chr11:3,795,000-8,504,999 bp) for cb-Ery (purple), nCD4 (orange), and Mon (pink) as calculated in each model ensembles. Error bar, indicating one standard deviation, is displayed for particles enclosing a transcribed gene (in at least one cell). The grey dashed line indicates 200 nm cut-off used in the analysis (**Methods**). **(B)** Cell-type specific distance distribution between particles enclosing the HBG2 gene and all transcribed genes in the genomic region (chr11:3,795,000-8,504,999 bp) for cb-Ery (purple), nCD4 (orange), and Mon (pink) as calculated in each model ensembles. Box boundaries represent 1st and 3rd quartiles, middle line represents median, and whiskers extend to 1.5 times the interquartile range (two-samples Kolmogorov-Smirnov test, asterisk indicate p-values < 7.5e^-6^). **(C-E)** Hierarchical clustering of each genes based on the co-occurrence analysis (**Methods**) in cb-Ery (**C**), nCD4 (**D**) and Mon (**E**). Co-occurrence value range from 0 (low, dark blue) to 100 (high, bright yellow). In each hierarchical tree the communities are labelled at their root branch. Per each gene the relative expression (log(FPKM) is shown in a scale of reds from 0 to 5. **(F)** Hierarchical clustering of the distances between the communities defined in cb-Ery (**Methods**). Distance values are coloured in the matrix from dark blue to bright yellow and the average expression in log(FPKM) per community is coloured by ranking from lowest (lightest) to highest (darkest) in 3 different shades of red. **(G)** Relationship between gene expression in log(FPKM) and the median distance of the gene particles to the centre of mass of its own community in cb-Ery ensemble of models (**Methods**). Purple line denotes the linear regression fit, the shading around the regression line represents the confidence interval, each community is represented with different symbols (circle community 1; inverse triangle community 2; square community 3; and ex community 4).

To further characterise the cell-type specific spatial distribution of these transcribed loci, we clustered their relative distances within the ensembles of 3D models and identified communities of expressed genomic loci (**Figure 4C-E** and **Methods**). Then, we quantified the amount of times a given community of expressed genomic loci occurred within the ensembles of 3D models (*i.e*., the co-occurrence score, **Methods**) and used this quantification as a proxy to define the “community stability”. This analysis revealed the existence of highly variable communities of expressed genomic loci that followed a cell-type specific segregation in the 3D space. Interestingly, the organization of these communities was overall more stable in cb-Ery than in nCD4 and Mon, where less defined communities were identified. Indeed, as assessed by the mean inter-community co-occurrence scores (**Methods**), the cb-Ery network was characterised by the presence of four stable communities (**Methods** and **Table 1**). While, the nCD4 network was formed by three communities with overall low co-occurrence (although community 2 in this network showed a stability in line with the communities in the cb-Ery network), and the Mon network formed by only two unstable communities (**Methods** and **Table 1**). Overall, the results highlight the presence of more defined 3D communities of expressed genes in cb-Ery as compared to nCD4 and Mon, suggesting that the co-occurrence of these segregated communities within an ensemble of possible folds is part of the cell-type specific 3D signature.

**Table 1.**
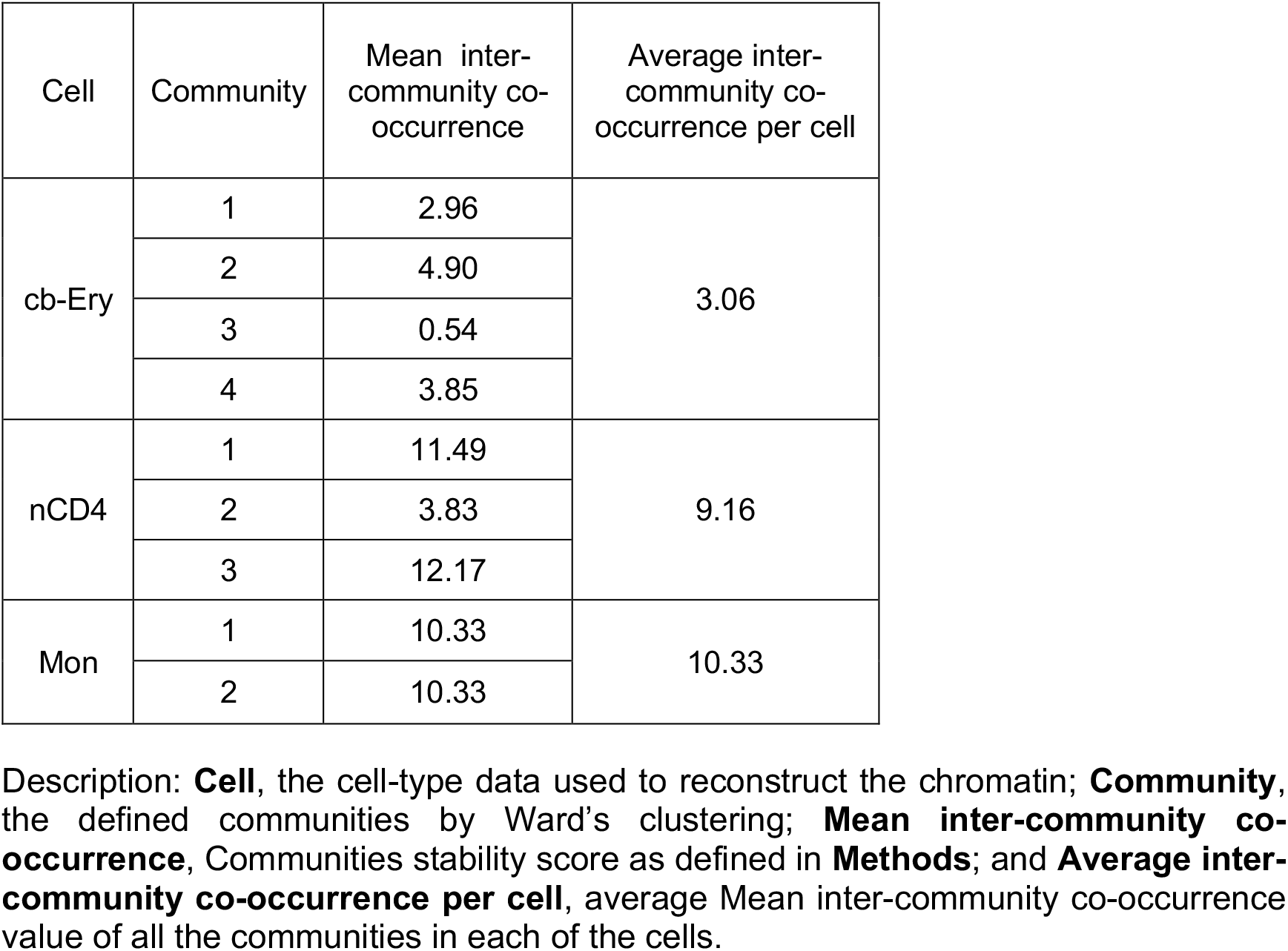
Communities stability assessment.

Next, we investigated whether the stability of the 3D communities of expressed genes in cb-Ery could be related to the high levels of expression of the β-globin genes (highest as HBG2 with 10.86 FPKM, while the mean expression of all the other expressed genes in nCD4 and Mon was 2.45 and 2.10 FPKM respectively). Clustering the distance distribution between the centres of mass of each community in cb-Ery (**Figure 4F**) revealed a clear hierarchical organisation with the most expressed community, which included the highly expressed β-globin locus (**Supplementary Table 2**), located in the centre, and the least expressed community in the periphery. This pattern was not present in nCD4, and impossible to address in Mon with just two communities (**Supplementary Figure 4A-B**). This suggests a hierarchical organisation in cb-Ery, in which the location in space of each of the communities and their levels of expression are related. Surprisingly, this hierarchy was also overall present at the community level in cb-Ery, where the distance between each gene to the centre of mass of the community and its expression were negatively correlated (CC: −0.55, p-value=0.002; **Figure 4G**). This suggests the formation in cb-Ery of a gradient of expression within the community were the most expressed genes are located in the centre of their communities and the less expressed ones are preferentially located in the periphery in line with the organisation previously observed for the alpha-globin locus [24]. This overall community organisation was not evident in nCD4 and Mon (**Supplementary Figure 4C-D**), thus suggesting that the high expression of the β-globin loci in cb-Ery could be associated with the establishment of a hierarchical organisation in the loci.

## Discussion

Here, we have introduced an integrative modelling method for the 3D reconstruction, analysis, and interpretation of sparse 3C-based datasets such as pcHi-C. We also demonstrate its usability in the comparative 3D analysis of genomic regions using the β-globin locus as an example, showing that our method can detect cell-type-specific 3D organisational features within genomic regions that can lead to several important implications on the relationship between genomic function and spatial genome organisation, such as the expression dependent organisation of active loci.

Generally, the analysis and interpretation of sparse 3C-datasets is not trivial and specialised analytical tools are required. In the case of pcHi-C, the available tools (ChiCMaxima, Chicago, Gothic, Chicdiff, HiCapTools [47-51]) are mainly focused on the implementation of normalization strategies to reduce the impact of non-biological biases and on strategies to detect interaction between captured loci. Conversely, the integrative modelling method presented in this study has been designed for the analysis and interpretation of sparse 3C-datasets in their third dimension, allowing for data normalisation, detection of significative interaction, and most importantly, the recovery of the full structural organization of a genomic region despite of the data sparseness.

Indeed, here we extensively tested our procedure by comparing models reconstructed directly from sparse and dense datasets, showing that 3D models reconstructed by the integrative modelling method for sparse data modelling are a good representation of the dense experiment. In fact, model reconstruction is only minorly effected by the intrinsic experimental biases of the capture experiment. Additionally, and most importantly, our model procedure reproduces remarkably well the whole 3D organisation of the selected genomic regions even recovering the organisation of loci that are not included as input restrains and are not readily observable in the sparse experiment.

Next, to assess whether the 3D reconstructed models were not only a bona fide representation of models based on Hi-C datasets, we used a ‘synthetic’ toy genome with known 3D organisation [38] and proved that we can efficiently model sparse pcHi-C-like datasets using as few as 2-3% of all possible interaction data. Importantly, this quantification highlights how the degree of sparseness of the data is related to the efficiency of the 3D reconstruction process and provide a general guideline for sparse data modelling. In light of this, we speculate that our integrative approach could easily be applied to different type of 3C datasets with similar sparseness. For example, protein-centric chromatin conformation method such as HiChIP [19] could be used as input experiment to reconstruct the chromatin folding, assuming that the protein-capture biases of this type of experiments are similar to the promoter-capture biases observed in the pcHiC experiments.

Finally, to illustrate the utility of our integrative approach, we applied it to the β-globin locus, whose 3D organisation has been extensively studied [39, 41, 52-54]. We investigated this locus in three different cell types (cb-Ery, nCD4, and Mon) and performed a comparative analysis between them. In agreement with previous studies [33], our models show that the topology of the β-globin locus varies in the three cell types owing to their differential expression. Interestingly, our models also unveil that the globin HBG2 gene is embedded in an epigenetically active and highly transcribed neighbourhood in cb-Ery giving rise to a locus-specific 3D functional signature. This functional signature is absent in the models of other cell-types (nCD4 and Mon), where the locus is not expressed. We also show that this cell-specific organisation, not only occurs proximally to the β-globin genes but also involves loci located at longer genomic distances (more than 1 Mb away). Indeed, our 3D comparative analysis unveiled the existence of an intricate cell-type specific network of spatially-proximal expressed genes that forms gene communities that are segregated in the 3D space in a cell-type specific fashion. The identified communities are compatible with the formation of chromatin foci in which transcribed genes co-localize as a general mechanism to organise gene transcription [24, 43-46, 55]. Interestingly, we observed that the co-occurrence within the ensemble of models of the identified cell-type specific communities is cell-type dependent, with the cb-Ery communities network formed by more persistent communities than the nCD4 and Mon community networks. This suggests that also the degree of co-occurrence of the communities within the ensemble is an important feature for the identification of a cell-type specific 3D signature. Additionally, we observed that in cb-Ery, where the β-globin genes are highly expressed, the communities present an overall hierarchical spatial organisation, both between and within communities. This topology is dependent on the level of transcription with highly expressed entities (entire community or specific gene within a community) located in the core of the hierarchical 3D organisation and low expressed entities found at the periphery. We hypothesise that the observed communities could represent cell-type specific transcription factories [24, 55-57] or phase-separated foci [58-60] organised following a gradient of transcription with high concentration of nascent transcripts and transcription machinery in the core of the assemblies that create a “sticky” environment to the less expressed peripheral loci. This hierarchical organisation is only marginally present in nCD4 and Mon, suggesting that it also contributes to the cell-type specific 3D signature characterising the β-globin region in cb-Ery.

In summary, we have shown that sparse datasets like pcHi-C can be effectively used to model in 3D the spatial conformation of genomic domains. The resulting models retain most of the genomic region organization and recover also the organisation of loci that are not readily observable in the sparse experiment. Importantly, this is achievable with a very small percentage (∼2-3%) of all possible interaction data in the genomic region. Additionally, our study not only provides a novel approach for sparse-data 3D modelling but also introduces new tools for the comparative analysis of genomic regions. Thus, it will aid the discovery of cell-type specific 3D signatures and help deciphering complex mechanism underlying the cell-type specific 3D genome organization.

## Methods

### Experimental datasets

Structural data were obtained from publicly available 3C-based chromatin interaction experiments of GM12878 cells (Hi-C GEO: GSE63525 and pcHi-C ArrayExpress: E-MTAB-2323) [6, 32], and cb-Ery, nCD4, and Mon cells (pcHi-C EGA: EGAS00001001911) [33].

#### Hi-C datasets processing

The reads for each replicate were mapped onto the GRCh38 reference genome, filtered, and merged using TADbit with default parameters [27]. Then, starting from the merged filtered fragments, the genome-wide raw interaction maps were binned at 5 kilo-base (kb) and normalized using OneD [37] as implemented in TADbit [27].

#### pcHi-C datasets processing

For each experiment, the reads were mapped onto the GRCh38 reference genome using TADbit [27] and were filtered applying the following filters: (i) self-circles, (ii) dangling-ends, (iii) errors, (iv) extra dangling-ends, (v) duplicated reads, and (vi) random breaks. Next, we computed the reproducibility score to measure the similarity between replicates from each pcHi-C dataset [61]. Then, for each cell-type, the different replicates from the same experiment were merged into one dataset for further analysis, making an exception with replicate ERR436029 from the GM12878 pcHi-C dataset (E-MTAB-2323), which was discarded due to a clearly low reproducibility score when compared with the rest of the replicates (average of 0.24 with the other replicates as compared to the average of 0.84 obtained between the other replicates). Using the merged filtered fragments, the genome-wide raw interaction maps of each cell-type were binned at 5 kb and normalised using the PRoportion of INTeraction approach (PRINT, next section).

#### Sparse data normalization

*PRoportion of INTeraction approach (PRINT)*. PRINT, a multi-stage normalisation procedure, weighs each pair of interacting bins with the same philosophy as the visibility approach for Hi-C [62]. Starting from a raw interaction matrix as input, PRINT first transforms the raw interaction between two bins (*i* and *j*) into a percentage of interaction with respect to the rest of the genome as:

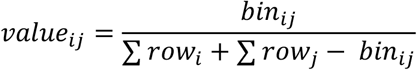

where (bin_*ij*_) represent the number of times in which bin *i* and *j* interact, and Σrow_*i*_ and Σrow_*j*_ are the sum of all the interactions of bins *i* and *j* respectively with all the genome (self-interactions included). Then, the non-baited interactions (that is, those bins containing only pcHi-C off-target reads) are filtered out.

#### PRINT assessment

Using the *benchmarking datasets* described above, each stage of PRINT normalisation (pcHi-C-raw, pcHi-C-pre and pcHi-C-norm) was assessed in comparison with the dense Hi-C interaction matrix by calculating the Spearman’s rank correlation coefficient between interactions (bin_*ij*_) present in both interaction matrices.

### Reconstructed 3D genomic regions

#### Benchmarking datasets

We selected 12 genomic regions of interest (**Supplementary Table 1**) as defined by Rao and colleagues [6]. This set of genomic regions were predicted to result in reliable 3D models based on their > 0.7 MMP scores [38] (**Supplementary Table 3**). Briefly, MMP score takes into account the interaction matrix size, the contribution of significant eigenvectors in the matrix, and the skewness and kurtosis of the z-scores distribution of the matrix to assess their potential for being modelled [38].

#### Comparative analysis datasets

We selected a genomic region around a locus of interest (here the β-globin) defining it in a semi-automatic manner in each cell type. Briefly, a viewpoint, which may be constituted by a bin or a set of bins of interest, is selected. Here, as viewpoint we used bins enclosing the active haemoglobin genes in cb-Ery (HBB, HBD, HBG1, and HBG2). Then, all the other bins that interacted with the viewpoint bins in the normalised genome-wide interaction matrix were selected. Each of these bins were then scored by their cumulative normalised interaction frequency values with the viewpoint bins. From this set only the top intra-chromosomal 200 bins were selected since, by visual inspection, they were the bins spanning the genomic region that best enclosed the viewpoint. Then an unweighted interaction network was generated with the nodes corresponding to the top 200 bins and the viewpoint bins. Edges between nodes were added if their pairwise cumulative normalised interaction frequency value was in the top 200 interacting bins. Then, a series of transformations were applied to the unweighted interaction network: (i) nodes that are highly proximal in 1D genomic resolution (closer than 25 kb) were merged into one node; and (ii) poorly connected nodes in the network that had less than 5 edges were filtered out (average number of edges per node in Mon, nCD4, and cb-Ery were 200, 214, and 214, respectively). The extreme nodes in terms of genomic coordinates were selected from the final unweighted interaction network to represent the optimal genomic region around the viewpoint. Here, to perform comparative analysis, we defined the optimal genomic region around the viewpoint as the broader genomic region that enclosed all of the genomic coordinates identified in each cell-type.

### 3D chromosome ensemble reconstruction from sparse datasets

#### Model representation

Each genomic region was described with a beads-on-string model based-on the previously implemented protocols [29, 63] without bending rigidity potential. Thus, a chromosome was represented with *N* spherical beads with diameter σ = 50 nm that contain 5 kb of chromatin which determined the genomic unit length of each model.

#### System set up for molecular dynamics

All simulations were done using TADdyn [29]. A generic random self-avoiding walk algorithm was used to define the initial conformation of each model. The potential energy of each system comprised the terms of the Kremer-and-Grest polymer model [64] including chain-connectivity (Finitely Extensible Nonlinear Elastic, FENE) [65] and excluded volume (purely repulsive Lennard-Jones) interactions. The initial conformation was placed randomly inside a cubic simulation box of size 1,000 σ centred at the origin of the Cartesian axis O = (0.0, 0.0, 0.0), tethered at the centre of the box using a harmonic (K_t_=50.0 k_B_T/σ^2^ and d_eq_=0.0 σ) to avoid any border effect and energy minimized using a short run of the Polak-Ribiere version of the conjugate gradient algorithm [66] to favour smooth adaptations of the implementations of the excluded volume and chain connectivity interaction.

#### Encoding sparse data into TADdyn restraints

TADdyn [29] empirically identifies the three optimal parameters to be used for modelling based on a grid search approach. This are: (1) maximal distance between two non-interacting particles (*maxdist*); (2) a lower-bound cut-off to define particles that do not frequently interact (*lowfreq*); and (3) an upper-bound cut-off to define particles that frequently interact (*upfreq*). All possible combinations of the parameters were explored in the intervals *lowfreq* = (−1.0,-0.5, 0, 0.5), *upfreq* = (−1, −0.5, 0, 0.5), *maxdist* = (200, 300, 400, 500) nm, and assessing each combination using distance thresholds to determine if two particles are in contact (*dcutoff*) at 100,150, 200, 250, 300, 350, 450, 500 nm. For each of the combinations an ensemble of 100 3D models was generated and the Spearman correlation coefficient between the contact map derived from each ensemble and the experimental input interaction matrix was calculated. The top set of parameters for each region in each cell-type were set for those resulting in the highest Spearman correlation coefficient between the models contact map and the input interaction matrix. To allow for a robust comparative analysis (**Methods**) the optimal *maxdist* and the *dcutoff* parameters were selected based on the consensus within the top optimal values for each region in each cell-type. Optimal *maxdist* and the *dcutoff* were set at 300 nm and 200 nm, respectively for the ensembles of models reconstructed from the GM12878, cb-Ery, nCD4, and Mon pcHi-C datasets. Once the three optimal parameters were defined, the type of restraints between each pair of particles was set considering an inverse relationship between the frequencies of interactions of the contact map and the corresponding spatial distances. Non-consecutive particles with contact frequencies above the upper-bound cut-off were restrained by a harmonic oscillator at an equilibrium distance, while those below the lower-bound cut-off were maintained further apart than an equilibrium distance by a lower-bound harmonic oscillator. To identify 3D models that best satisfy all the imposed restraints, the optimization procedure was then performed using a steered molecular dynamic protocol. A total of 1,000 replicate trajectories were generated for each genomic region and dataset. Per each of the 1,000 replicate trajectories, the conformation at the end of the steering protocol (when the target spring constant and equilibrium distance are reached) was retained to form the final ensemble of 1,000 3D models. For the cb-Ery, nCD4, and Mon datasets, to account for possible mirrored 3D models within the final ensemble of 3D models, each ensemble was then clustered based on structural similarity score as implemented in TADbit [27] and only the models from the most populated cluster were retained for further analysis.

#### Steered Molecular Dynamics protocol

A steered molecular dynamics protocol was used to progressively favour the imposition of the defined set of restrains between non-consecutive particles. For each restraint, the equilibrium distance was set to 1 particle diameter (σ). The spring constant *k(L,t)* was weighted with the sequence-separation *L* between the constrained beads as in TADdyn [29] to ensure that the steering process was not dominated by the target pairs at the largest sequence separation. However, here the *k(L,t)* was smoothly ramped during the steering phase from zero to its maximum value.

### 3D chromosome ensemble reconstruction from dense datasets

The reconstruction of 3D models of genomic regions from dense data followed the modelling protocol described above. That is, a grid search approach was used to select for the optimal parameters to be used for modelling. The optimal *maxdist* and the *dcutoff* parameters were selected based on the consensus within the top optimal values for each region in the GM12878 pcHi-C dataset and set at 300 and 200 nm, respectively. Using these parameters, the final ensemble of 1,000 3D models was obtained starting from the computed 1,000 steered molecular dynamics trajectories.

### 3D chromosome ensemble reconstruction from Virtual pcHi-C derived from dense datasets

A dataset of Virtual pcHi-C interaction matrices was produced starting from the normalised Hi-C interaction matrices at 5kb resolution (GM12878 cells GEO: GSE63525; **Methods**) and from the liftover (https://genome.ucsc.edu/cgi-bin/hgLiftOver) list of captured fragments in pcHi-C GM12878 experiment [32]. The obtained Virtual pcHi-C interaction matrices comprised only interactions (bin_*ij*_) in which either *i* or *j* enclose the coordinates of a captured fragment. These interaction matrices were used as input for the reconstruction of 3D models of genomic regions following the modelling protocol described above. The optimal *maxdist* and the *dcutoff* parameters were set at 300 and 200 based on their consensus with the parameters used in the GM12878 pcHi-C dataset. A total of 1,000 steered molecular dynamics trajectories were computed, and for each trajectory the conformations satisfying the majority of the imposed constraints within a radius of 2 σ were retained.

### 3D chromosome ensemble reconstruction from ‘synthetic’ sparse dataset

We used a previously published “toy genome” [38] (that is, the ensemble of models accounting for the formation of TAD-like architecture with low structural variability and high noise levels that comprises a total of 626 particles at the highest genomic resolution) to randomly select 10 sets of 22 loci from the toy genome contact map (or synthetic interaction maps). These loci mimic pcHi-C to generate reliable sparse interaction matrices comprising only interactions (bin_*ij*_) in which either *i* or *j* have been selected as random captured loci. Each of these sets was then randomly subsampled to generate ‘synthetic’ capture matrices with 2, 4, 6, 10, 14, and 18 selected captured loci. The obtained ‘synthetic’ capture matrices (70 in total) were next used as input for the reconstruction of 3D models of genomic regions following the modelling protocol described above. The optimal *maxdist* and the *dcutoff* parameters were set at 500 and 200 nm. Using these parameters, a final ensemble of 100 3D models was reconstructed for each ‘synthetic’ capture matrices comprising the conformations that best satisfied the imposed restraints in each of the computed 100 steered molecular dynamics trajectories.

### Analysis of the ensemble of 3D models

#### Contact map generation

For each ensemble of 3D models, a contact map was calculated at 5kb resolution to visualize the frequencies of contacts in the ensemble. Two beads were considered to constitute a contact when their Euclidean distance was below 200 nm cut-off.

#### Matrix Comparison

The degree of similarity between two matrices was computed by comparing each cell from the matrices, or a subset of them, using the Spearman’s rank correlation coefficient (*r*_*s*_) as implemented in the Python library SciPy [67, 68]:

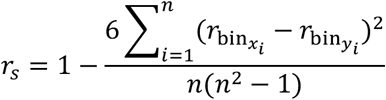

where 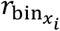 is the rank of the *i*^*th*^ observation in one matrix, 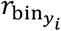 is the rank of the *i*^*th*^ observation in the other matrix, and *n* states for the number of pairs of observations.

#### Particle-to-particle median distance correlation (ppMdC)

For each ensemble of 3D models, we differentiated 3 sets comprising particles enclosing the coordinates of: (i) captured loci (capture), (ii) non-captured loci (other), and (iii) all the loci (all). For each of the pairs of particles in a given set we calculated the particle-to-particle median distance. Then, the degree of similarity between two given sets was computed using the Spearman’s rank correlation coefficient between their particle-to-particle median distances. The ppMdC measure varies between −1.0 and 1.0 for comparisons where the particle-to-particle median distances perfectly anti-correlate or correlate, respectively.

#### Hierarchical clustering of ensembles of 3D models

Multiple ensembles of 3D models were merged in a unique set and the models were structurally superpose using pair-wise rigid-body superposition. Next, the all-vs-all distance root mean square deviation (dRMSD) was calculated and the resulting dRMSD matrix was hierarchically clustered using Ward’s sum of squares method [69] as implemented in the Python library SciPy [67].

#### Cell-specific expression profile

Publicly available [33] expression matrix containing the expression values (log(FPKM)) of each gene in cb-Ery, nCD4, and Mon cell types was downloaded (GeneExpressionMatrix.txt.gz at https://osf.io/u8tzp/). The 3 datasets had two or more replicates each (2 cb-Ery, 5 Mac, and 8 nCD4, respectively), thus the average expression value of each gene from all replicates was used. Then, a cell-specific per-bin cumulative expression profile of the chr11:3,795,000-8,505,000 genomic region at 5kb resolution was obtained assigning the mean expression value of each gene (with log(FPKM)>0) to bins enclosing for the coordinates of its transcription start site (coordinates retrieved from bioMart [70]).

#### 3D enrichment analysis

To study the spatial co-localization of different regulatory elements and the local levels of transcription (based on genome-wide ChIP- and RNA-seq data) around a selected locus (central viewpoint) we implement a *3D enrichment analysis tool* (named ‘radial-plot’) that allows the comparison of heterogeneous sets of data from multiple data sources. Per each cell type a per-particle binarized chromatin marks profile in the genomic region was generated starting from the ChIP-seq signal of H3K27ac, H3K36me3, H3K4me1, H3K4me3, H3K9me3, and H3K27me3 in cb-Ery, nCD4, and Mon cell types [33]. A particle was considered enclosing for a chromatin mark if a peak was present. Similarly, we also constructed, for each cell type, a per-particle binarized transcription profile starting from the cell-specific expression profile (**Methods**). Then the 3D spatial distribution of the 3D enrichment based on the per-particle binarized profile around the chosen central viewpoint was calculated as follow: (i) starting from the central viewpoint an initial sphere with a radius of 200 nm was constructed; (ii) a series of spherical shells, that occupied a volume equal the initial sphere, were added; (iii) per each model in the ensemble of 3D models a particle of the binarized profile was assigned to a spherical shell based on its relative distance to the central viewpoint; (iv) per each spherical shell we performed Fisher’s exact tests for 2 × 2 contingency tables comparing the amount of particles with or without signal in the spherical shell with the outside ones, and the log of the odd ratios was assigned to the shell if the p-value < 0.01. The obtained 3D enrichment was then visualised as a 2D radial plot.

#### Defining gene communities

*co-occurrence of expressed genes*. For each ensemble of 3D models, based on their cell-specific expression profile (**Methods**), we defined the set of expressed particles (log(FPKM) > 0). Then, considering this set of particles, an all-vs-all pairwise distances matrix was calculated in each model and hierarchically clustered using Ward’s sum of squares method [69] as implemented in the Python library SciPy [67]. Then the Calinski-Harabasz index [71], as implemented in the Python library Scikit-learn [72], was used to determinate the optimal number of clusters in each dendrogram. Then, for each ensemble, a co-occurrence matrix was generated considering the percentage of models in which a pair of particles belonged to the same cluster. The co-occurrence measure varies between 0 and 100, where 0 indicates absence of co-occurrence and 100 indicates a stable co-occurrence within the ensemble of 3D models. The co-occurrence matrix was next hierarchically clustered using Ward’s sum of squares method [69] and communities of co-occurrent active genes were identified using the Calinski-Harabasz index analysis in the dendrogram.

#### Communities stability within the ensemble of models

To assess the stability of each community within the ensemble we introduced the inter-community co-occurrence score that defines the degree of unstable compositions of a community. It is computed as the mean co-occurrence values between each gene in a community and the rest of the communities.

#### Distance between communities and within community

To describe the spatial arrangement of each community for a given ensemble of 3D models, we treated each community as a rigid body and calculated its centre of mass (COM) in each 3D model of the ensemble. Per each model the all-vs-all pairwise distances between the COMs of each communities were computed and the mean distance values assigned as the typical distance between communities. Similarly, per each model, we also calculated the distance of each particle in a given community and the COM of its community. The within community distance of a given particle was defined by its mean value in the ensemble of 3D models.

## AUTHOR CONTRIBUTIONS

JM-E, IF and MAM-R conceived the study; JM-E, MDS, and IF performed the modeling; DC supported modeling protocol development and implementation; JM-E, IF wrote the manuscript with MDS, DC and MAM-R; IF and MAM-R oversaw the project.

## ACKNOWLEDGMENTS

We thank all the current and past members of the Marti-Renom lab for their continuous discussions and support. Dr. Irene Miguel-Escalada for helpful discussions. The 4D genome unit at CRG for data availability. Javierre Lab for providing access to the ChIP-seq peaks for the beta-globin locus in different cell types. We acknowledge the ENCODE consortium and the ENCODE production laboratories that generated the datasets used in the manuscript. This study makes use of data generated by the PCHI-C Consortium available in the EGA European Genome-Phenome Archive (National Institute for Health Research of England, UK Medical Research Council (MR/L007150/1) and UK Biotechnology and Biological Research Council (BB/J004480/1)).

