## Supplementary_Figures_Tables for "3D reconstruction of genomic regions from sparse interaction data"

### Supplementary Figures and Tables

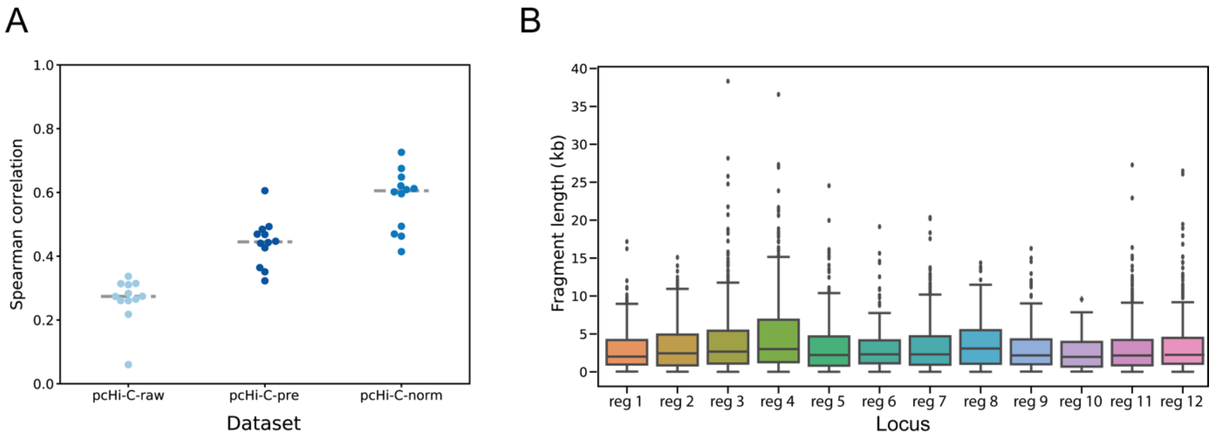


**Supplementary Figure 1. Integrative modelling procedure: assessing Print normalisation procedure and defining the model representation. (A)** Assessing Print multi-step procedure. Element-wise Spearman correlation between each stage of Print normalisation (pcHi-C-raw in light blue; pcHi-C-pre in dark blue; and pcHi-C-norm in medium blue) and the Hi-C interaction matrix. The grey dashed line indicates the median correlation in the entire benchmark dataset at each stage. **(B)** Distribution of the sizes (in kb) of the restriction fragments in each of the regions comprised in the benchmark dataset.

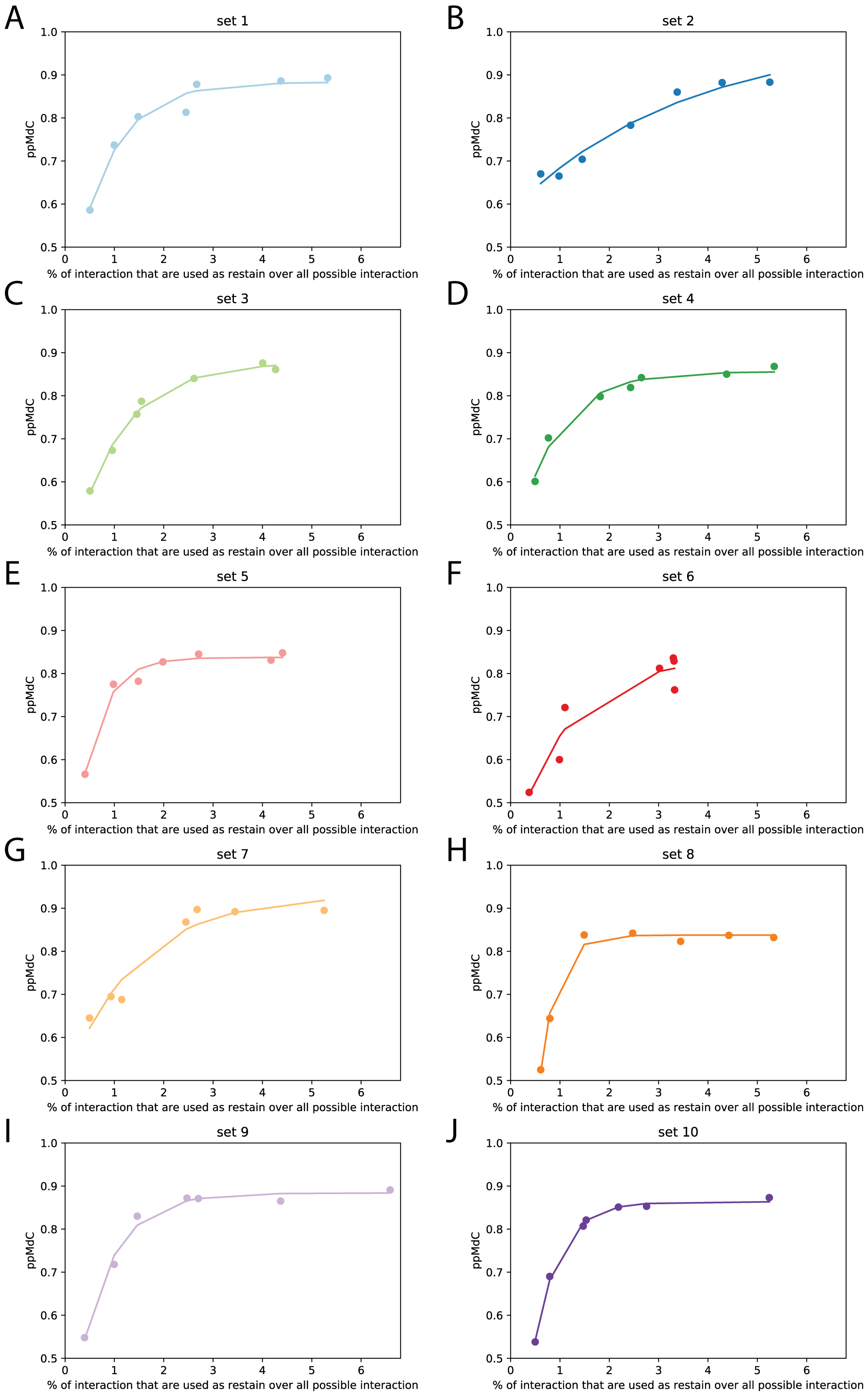
A
set 1
B
set 2
C
set 3
D
set 4
E
set 5
F
set 6
G
set 7
H
set 8
I
set 9
J
set 10
ppMdC
1.0
0.9
0.8
0.7
0.6
0.5
0
1
2
3
4
5
6
% of interaction that are used as restain over all possible interaction

**Supplementary Figure 2. Comparison of each 'synthetic' capture sets and the toy genome. (A-J)** Relationship between the ppMdC and the percentage of cells in the matrix used as restrains in each set. The dots in the plot represent the degree of sparseness in each subset (2, 4, 6, 10, 14, 18, and 22 captures) and the coloured line indicates the fitted exponential function. The colour code used is the same as in Figure 2C.

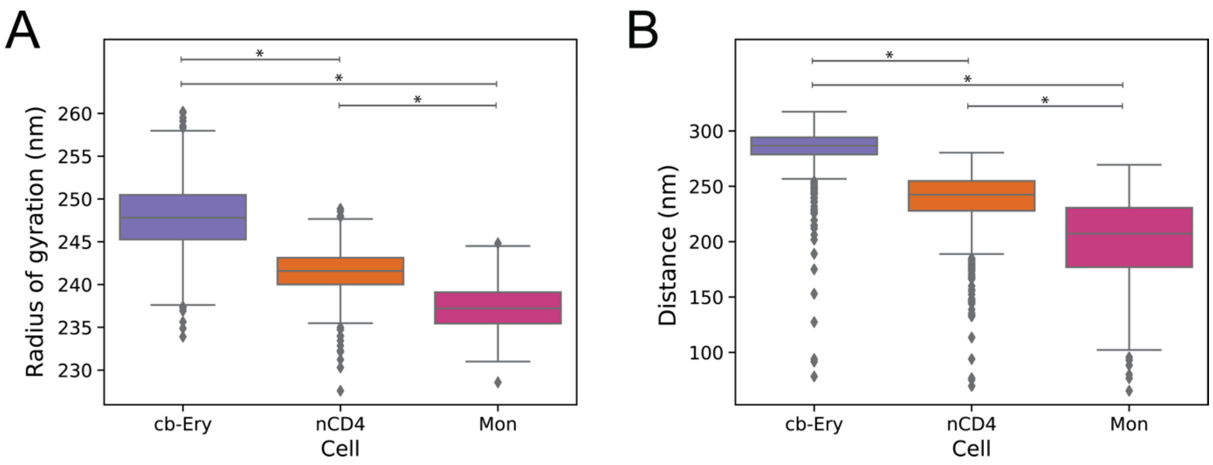


**Supplementary Figure 3. Cell-type-specific 3D features in the model ensembles. (A)** Cell-type specific distribution of the radius of gyration of the models in the ensemble. Box boundaries represent 1st and 3rd quartiles, middle line represents median, and whiskers extend to 1.5 times the interquartile range (two-samples Kolmogorov-Smirnov test, asterisk indicate $p < 9.1e^{-163}$). **(B)** Cell-type specific distance distribution of the centres of mass of the particles containing the β-globin genes (HBB, HBD, HBG1, HBG2, and HBE1) from the centre of the model as calculated in each model of the ensemble. Box boundaries represent 1st and 3rd quartiles, middle line represents median, and whiskers extend to 1.5 times the interquartile range (two-samples Kolmogorov-Smirnov test, asterisk indicate $p < 3.46e^{-101}$).

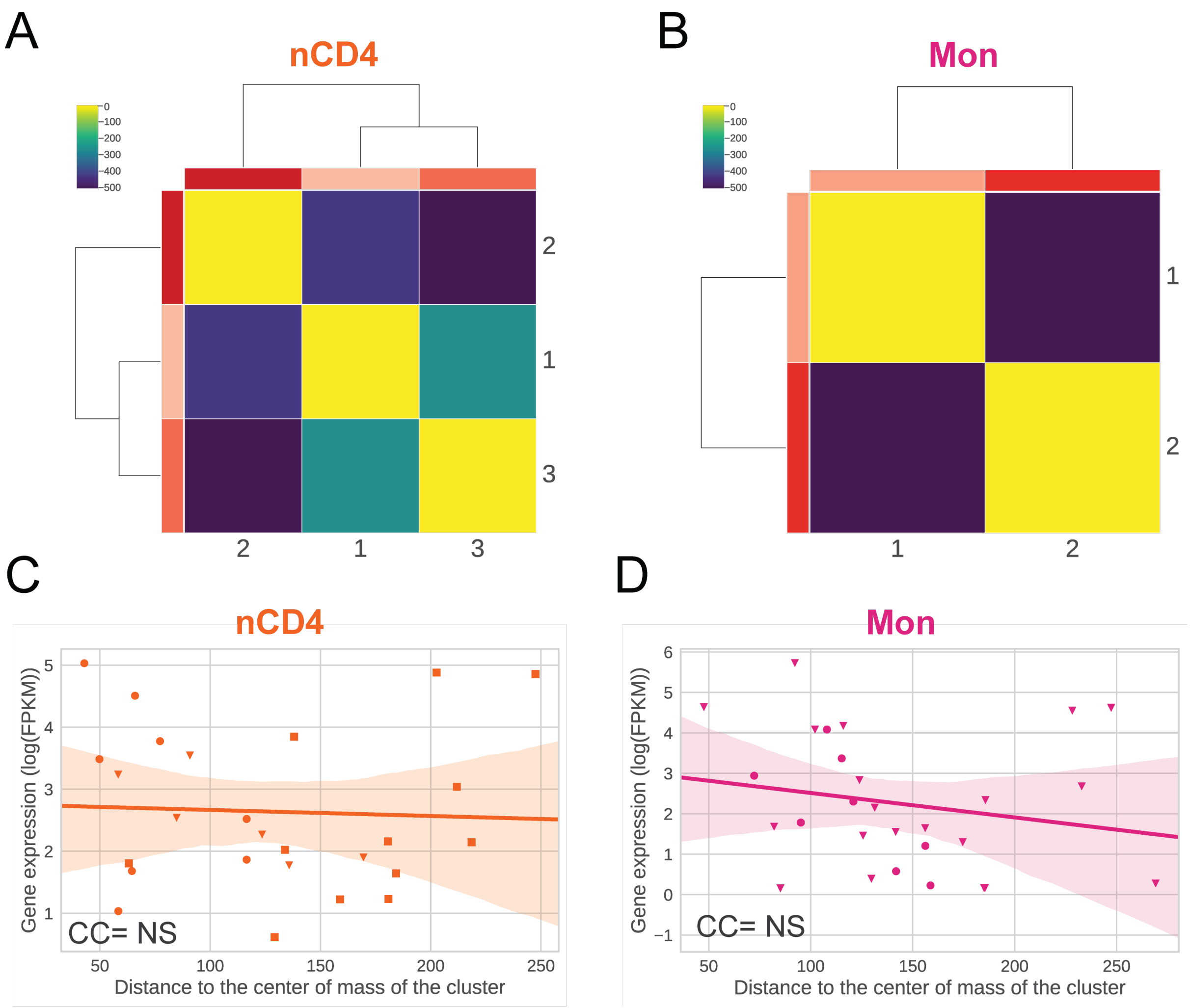


**Supplementary Figure 4. Hierarchical 3D organisation of expressed entities in nCD4 and Mon. (A-B)** Hierarchical clustering of the distances of mass (**Methods**) between the communities defined in nCD4 **(A)** and Mon **(B)**. Distance value is coloured in the matrix from dark blue (low) to bright yellow (high) and the average expression in log(FPKM) per community is coloured by ranking from lowest (lightest) to highest (darkest) in 3 **(A)** and 2 **(B)** different shades of red. **(C-D)** Relationship between gene expression in log(FPKM) and the median distance of the gene particles to the centre of mass of its own community in nCD4 **(C)** and Mon **(D)** ensemble of models (**Methods**). Orange **(C)** and pink **(D)** line denote the linear regression fit, the shading around the regression line represents the confidence interval, each community is represented with different symbols (circle community 1; inverse triangle community 2; square community 3; and ex community 4) ; NS stand for not significant.

**Supplementary Table 1**. Benchmark datasets

| **Locus** | **Chromosome** | **Start** | **End** |
|---|---|---|---|
| Region 1 | chr7 | 137,515,000 | 138,120,000 |
| Region 2 | chr8 | 132,755,000 | 133,560,000 |
| Region 3 | chr20 | 50,745,000 | 52,515,000 |
| Region 4 | chr3 | 49,325,000 | 51,095,000 |
| Region 5 | chr3 | 63,110,000 | 64,715,000 |
| Region 6 | chr8 | 119,785,000 | 120,190,000 |
| Region 7 | chr17 | 68,500,000 | 70,005,000 |
| Region 8 | chr2 | 10,705,000 | 11,210,000 |
| Region 9 | chr10 | 88,890,000 | 89,495,000 |
| Region 10 | chr1 | 169,590,000 | 169,745,000 |
| Region 11 | chr21 | 26,625,000 | 28,930,000 |
| Region 12 | chr13 | 84,575,000 | 86,180,000 |

Description: **Locus**, the name of the region modelled starting from Hi-C, pcHi-C, and pcHi-Cvirt datasets; **Chromosome**, the chromosome where the region is located; **Start** and **End**, represent the genomic coordinates (GRCh38 assembly).

**Supplementary Table 2**. Gene communities expression statistics

| Cell | Community | nGenes | MeanExp | σ |
|---|---|---|---|---|
| cb-Ery | 1 | 8 | 2.48 | 0.81 |
| | 2 | 11 | 4.57 | 3.28 |
| | 3 | 3 | 1.57 | 1.99 |
| | 4 | 6 | 2.78 | 0.78 |
| NCD4 | 1 | 8 | 2.48 | 0.81 |
| | 2 | 6 | 4.57 | 3.28 |
| | 3 | 12 | 1.57 | 1.99 |
| Mon | 1 | 8 | 2.48 | 0.81 |
| | 2 | 20 | 4.57 | 3.28 |

Description: **Cell**, source cell type data used for the modelling approach; **Community**, the number assigned to each community; **nGenes** , number of active genes composing each community; **MeanExp**, mean expression of the genes within the community; **σ**, standard deviation of the mean expression of genes within the community.

**Supplementary Table 3**. MMP scores of the 12 modelled Hi-C interaction matrices

| Dataset | Locus | MMP score |
|---|---|---|
| HiC data (GSE63525) | Region 1 | 0.80 |
| | Region 2 | 0.78 |
| | Region 3 | 0.76 |
| | Region 4 | 0.78 |
| | Region 5 | 0.78 |
| | Region 6 | 0.81 |
| | Region 7 | 0.78 |
| | Region 8 | 0.81 |
| | Region 9 | 0.82 |
| | Region 10 | 0.84 |
| | Region 11 | 0.72 |
| | Region 12 | 0.75 |

Description: **Dataset**, experimental dataset used to reconstruct the 12 ensembles of models; **Locus**, the name of the region modelled starting from the Hi-C dataset; **MMP score**, Value of the MMP score of the interaction matrix of each of the locus. It predicts the reliability of the 3D models based on the interaction matrix size, the contribution of significant eigenvectors in the matrix, and the skewness and kurtosis of the z-scores distribution of the matrix [1, 2].


1. Trussart M, Serra F, Bau D, Junier I, Serrano L, Marti-Renom MA. Assessing the limits of restraint-based 3D modeling of genomes and genomic domains. Nucleic Acids Res. 2015;43(7):3465-77. doi: 10.1093/nar/gkv221. PubMed Central PMCID: PMCPMC4402535.
2. Serra F, Bau D, Goodstadt M, Castillo D, Filion GJ, Marti-Renom MA. Automatic analysis and 3D-modelling of Hi-C data using TADbit reveals structural features of the fly chromatin colors. PLoS Comput Biol. 2017;13(7):e1005665. doi: 10.1371/journal.pcbi.1005665. PubMed Central PMCID: PMCPMC5540598.
